# Negative selection on a *SOD1* mutation limits canine degenerative myelopathy while avoiding inbreeding

**DOI:** 10.1101/2023.07.25.550492

**Authors:** Hisashi Ukawa, Noriyoshi Akiyama, Fumiko Yamamoto, Ken Ohashi, Genki Ishihara, Yuki Matsumoto

## Abstract

Several hundred disease-causing mutations are currently known in domestic dogs. Breeding management is therefore required to minimize their spread. Recently, genetic methods such as direct-to-consumer testing have gained popularity; however, their effects on dog populations are unclear. Here, we aimed to evaluate the influence of genetic testing on the frequency of mutations responsible for canine degenerative myelopathy (DM) and assess the changes in the genetic structure of a Pembroke Welsh corgi population from Japan. Genetic testing of 5,512 dogs for the causative mutation in superoxide dismutase 1 (*SOD1*) (c.118G>A (p.E40K)) uncovered a recent decrease in frequency, plummeting from 14.5% (95/657) in 2019 to 2.9% (24/820) in 2022. Weir and Cockerham population differentiation (*F*_ST_) and simulation-based genome-wide single-nucleotide polymorphism (SNP) analysis of 117 selected dogs revealed 143 candidate SNPs for selection. The SNP with the highest *F*_ST_ value was located in the intron of *SOD1* adjacent to the c.118G>A mutation, supporting a strong selection signature on *SOD1*. Further genome-wide SNP analyses revealed no obvious changes in inbreeding levels and genetic diversity between the 2019 and 2022 populations. Our study highlights that genetic testing can help inform improved mating choices in breeding programs to reduce the frequency of risk variants and avoid inbreeding. This combined strategy could decrease the genetic risk of canine DM, a fatal disease, within only a few years.

**Significance statement:** Genetic breeding methods using direct-to-consumer testing have gained popularity, but their effects on dog populations remain unclear. In this study, the effect of direct-to-consumer genetic testing on *SOD1* mutation, the causative element of canine degenerative myelopathy, in a domestic dog population (Pembroke Welsh corgi) from Japan was investigated. Our analyses revealed that since the expansion of genetic testing in 2019, breeders used these tests to artificially select against the *SOD1* mutation, considerably decreasing its occurrence in the corgi population within only a few years (2019 versus 2022). Our study makes a substantial contribution to existing literature by providing empirical evidence that direct-to-consumer genetic testing can have rapid influence on pet genetics, noticeable in a span of 2–3 years.

## Introduction

Genetic testing for disease-causing mutations in companion animals is increasingly performed by veterinarians for diagnosis, by breeders to reduce the incidence of inherited disease, and even by pet owners to determine the genetic background of their pets (Moses et al. 2018). Genetic tests employed by pet owners are termed direct-to-consumer (DTC) testing and can be broadly classified into two categories: 1) detection of specific mutations using sequencing or probes and 2) high-throughput genotyping using genome-wide marker sets designed to detect multiple mutations simultaneously. In addition to their convenience, these DTC tests also yield data with substantial implications for genetic research. For example, a recent study using DTC test samples revealed breed-specific genetic mutations associated with hypertrophic cardiomyopathy in several cat breeds (Akiyama et al. 2023). Another study, employing over 10,000 DTC samples, determined the allele frequencies of 12 genes associated with canine coat color and the physical characteristics of different dog lineages (Dreger et al. 2019). The results indicated that random mating between certain dog breeds can produce unexpected phenotypes, including embryonic lethality (Dreger et al. 2019). Moreover, although at least 775 disease-associated mutations are already known in dogs (Rokhsar et al. 2021), a recent genome-wide association study using DTC genetic testing data revealed a novel and unexpected in-frame deletion that causes deafness in Rhodesian ridgebacks (Kawakami et al. 2022). Thus, the widespread adoption of animal genetic testing greatly benefits genetic studies.

Genetic testing for adult dogs was introduced in Japan in 2017, followed by testing for puppies in 2019. The results of such widespread testing could affect dog populations by preventing carriers from breeding, thus limiting the number of genetically affected animals. However, the actual effect of widespread testing on mutation frequency remains unclear. In addition, while breeding to avoid mutant alleles could affect the genetic structure and inbreeding levels in dog populations, only a few studies have investigated this possibility.

Since the initial report of the draft genome of domestic dogs in 2005 (Lindblad-Toh et al. 2005), significant advancements have been made in canine genetic research. Inbreeding and genetic structures have been evaluated at the population level based on microsatellite and genome-wide single-nucleotide polymorphism (SNP) analyses (Boyko et al. 2009; Mellanby et al. 2013; Dreger et al. 2016; Chu et al. 2019). Accordingly, genome-wide SNP analyses can be employed to evaluate the influence of expanded DTC genetic testing on genetic structure in dog populations.

Canine degenerative myelopathy (DM) is a fatal neurodegenerative disease prevalent in several dog breeds, including the Pembroke Welsh corgi (PWC), German shepherd, and boxer (Neeves and Granger 2015). The c.118G>A mutation in the superoxide dismutase 1 (*SOD1*) gene (p.E40K on chromosome 31; 26,540,342 bp, based on CanFam 3.1) is reportedly a causative factor of DM in PWCs (Awano et al. 2009). SOD1 is one of the two antioxidant isozymes responsible for specifically eliminating free superoxide radicals in mammals. The homozygous A allele mutation is strongly associated with DM onset (Awano et al. 2009; Chang et al. 2013; Zeng et al. 2014), indicating that it is an autosomal recessive variant. The mutant homozygous A allele is fairly common in PWCs and is not geographically restricted (e.g., 48.4% or 59/122 in Japan (Chang et al. 2013) and 83% or 14/21 in Mexico (Ayala-Valdovinos and Gomez-Fernandez 2018)).

In this study, we evaluated the allelic frequency of *SOD1*: c.118G>A (p.E40K) mutation in a population of over 5,500 PWCs from Japan, analyzing DTC genetic testing data across three years (2019–2022). PWCs born from 2012 to 2022 were included in this dataset, allowing us to examine the impact of genetic testing on the PWC population for both adult dogs and puppies that were introduced in 2017 and 2019, respectively. We also performed a genome-wide analysis to detect and compare selection signatures between the populations in 2019 and 2022. Finally, we assessed inbreeding and population structures based on genome-wide SNPs to determine the effects of genetic testing on dog breeding. The findings of this study could provide valuable insights into how widespread genetic testing controls the spread of genetic disorders among dogs.

## Results

### Allele frequencies of the SOD1: c.118G>A variant

We examined the distribution of *SOD1*: c.118G<A by birth year in 5,512 PWCs (Fig. 1A). Most PWCs born between 2012 and 2017 were homozygous for the mutant allele (Mutant) or heterozygous (Hetero), whereas homozygous wild-type (Wild) exhibited the lowest frequency (9.0% or 12/134 in 2013 to 15.3% or 44/287 in 2016). With the initiation of genetic testing for adult dogs in 2017, the frequency of the Mutant decreased from 39.4% (163/414) in 2017 to 2.9% (24/820) in 2022.

**Figure 1.**
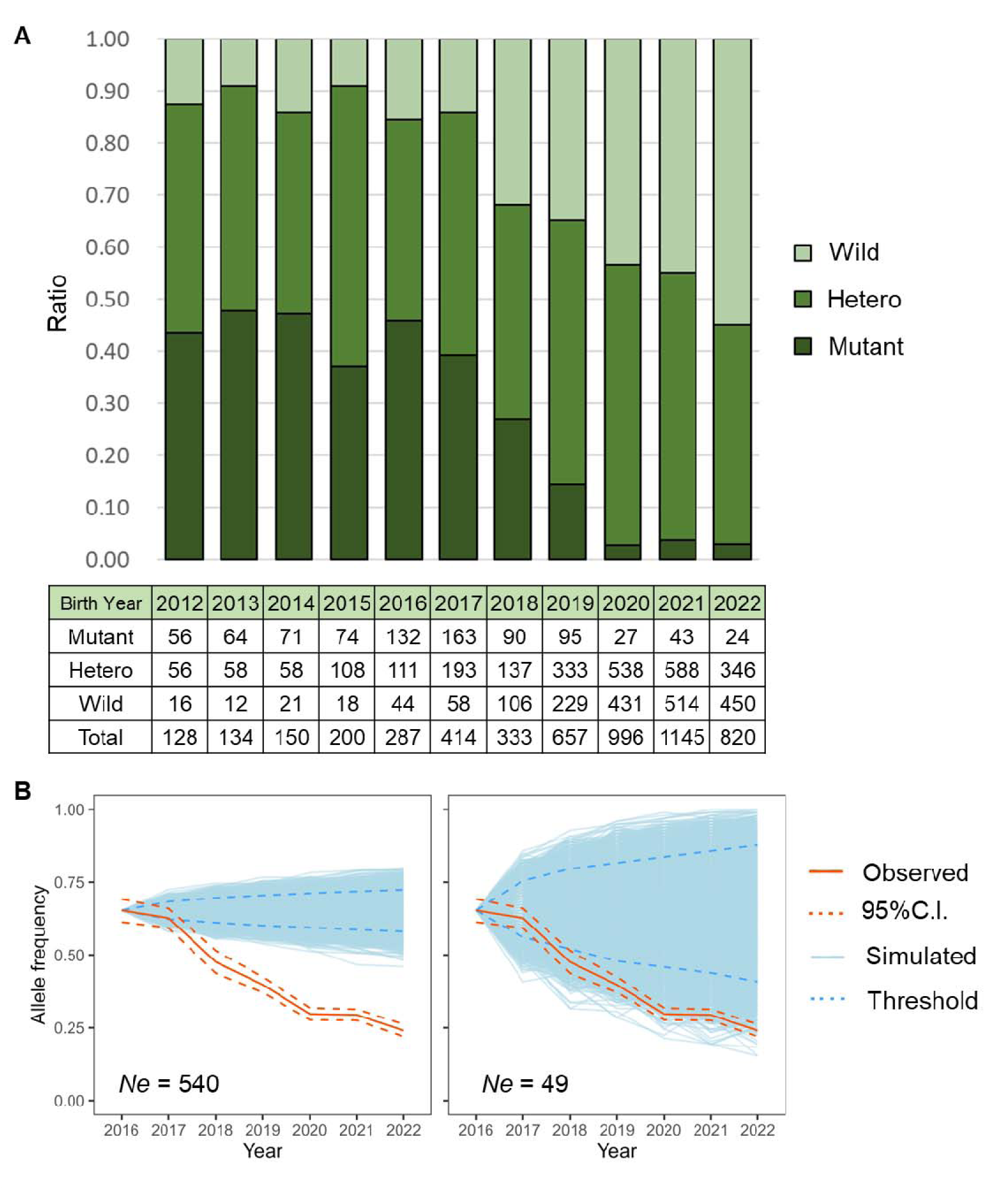
Trends in diploid genotype and allele frequencies of the *SOD1*: c.118G>A mutation. A: Diploid genotype frequencies of the *SOD1*: c.118G>A mutation from 2012 to 2022. The ratio of allele frequency is based on the birth year of tested dogs. The lower chart shows the number of tested dogs divided by each diploid genotype. B: Real and simulated allele frequencies of the *SOD1*: c.118G>A mutation for six years after 2016 based on two models: large (*Ne* = 540, left) and small (*Ne* = 49, right) models (see Methods for details). The orange line indicates the observed allele frequency, and the dashed orange lines indicate 95 percentile confidence intervals. The light-blue lines indicate 10,000 simulated allele frequencies starting from 2016, and the dashed blue lines indicate 95 percentile thresholds.

Since 2019, when genetic testing in puppies began, the ratio of each diploid genotype has changed significantly, considering the ratios observed in 2022 (Fisher’s exact test, *P* = 0.001). Specifically, the proportion of Mutants decreased from 14.5% (95/657) in 2019 to 2.9% (24/820) in 2022.

To further investigate the allele frequency of the *SOD1*: c.118G<A mutation, we performed a simulation based on random genetic drift (10,000 replicates) for two models (i.e., scenarios where effective population sizes (*Ne*), which refers to the number of bred dogs, were larger (*Ne* = 540) or smaller (*Ne* = 49); see Methods for details). Deviation from genetic drift would indicate selection. The observed allele frequency was significantly lower than the simulated one after 2018 in both models (*P* < 0.05; Fig. 1B, Table S1).

### Selection signature

To reveal the existence of selection signatures for *SOD1* and other regions at the genome-wide level, we compared the 2019 and 2022 groups to determine the selection signature in the PWC genome. Genome-wide SNP-based analyses on 117 PWC pups were performed after determining their genetic backgrounds and separating them into four groups according to the *SOD1*: c.118G<A genotype (Table S2). We set the Weir and Cockerham population differentiation (*F*_ST_) and simulation-based thresholds, and all SNPs with the top 0.1% *F*_ST_ values met the simulation-based thresholds (Table S3; see Methods for details). Using both thresholds, we selected 5 SNPs and identified 138 SNPs as candidates for selection (Table S3). The SNP “BICF2G630738971” had the highest *F*_ST_ (0.25) of the 143,013 tested SNPs, and this SNP was located in the intron of *SOD1* on canine chromosome 31 (Fig. 2A, B), 10,529 bp downstream of *SOD1*: c.118G>A. We then calculated the extended haplotype homozygosity (EHH) of each population (2019 vs. 2022) for BICF2G630738971 (Fig. 2C). The 2019 group had longer EHH haplotypes than the 2022 group, although the position of *SOD1*: c.118G>A was closely linked to the top SNP in the 2022 group (0.47 < EHH < 1).

**Figure 2.**
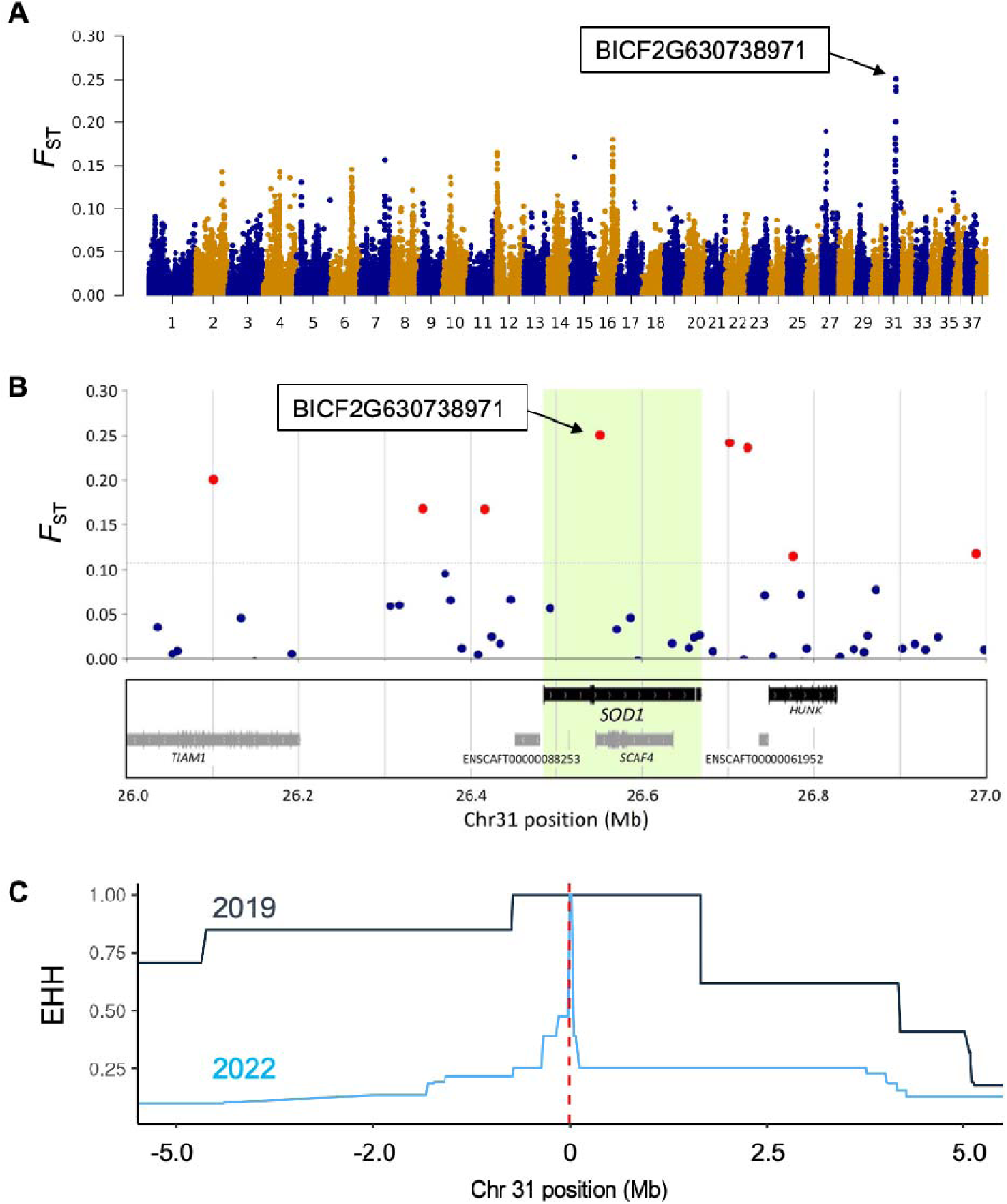
Selection signature observed in *SOD1*. A: Manhattan plot based on *F*_ST_. B: Relationships of *F*_ST_ per SNP and their locations around *SOD1* on chromosome 31. SNP BICF2G630738971 had the highest *F*_ST_ value. Red dots indicate SNPs that exceeded the threshold of 0.1% *F*_ST_. C: EHH of each group from the top *F*_ST_ SNP (BICF2G630738971) for the derived allele. Dark-blue and light-blue lines indicate the 2019 and 2022 groups, respectively. The red dotted line indicates the position of *SOD1*: c.118G.

To investigate any other genes under selective pressure in addition to *SOD1*, we surveyed the 186 genes that were included in or near the selected or candidate SNPs for selection (Table S4). Four protein-coding genes (*ABCA4*, *TNXB*, *COL11A2*, and *SOD1*) reported to affect phenotypes were identified (Fig. S1, Table S4). Gene Ontology (GO) analysis revealed three significantly enriched terms (Fig. S2 and Table S5); one was related to peptide antigen binding, and the others were related to the major histocompatibility complex (MHC).

### Inbreeding levels and genetic structure

We assessed inbreeding levels in the dog population. Observed heterozygosity (Ho) per group was calculated using 142,510 SNPs; the Ho was 0.305 and 0.306 for the Wild 2019 and Wild 2022 groups, respectively. For mutant PWCs, the Ho was 0.296 and 0.314 in 2019 and 2022, respectively. We then compared the inbreeding coefficients between groups based on runs of homozygosity (F_ROH_) (Fig. 3). The mean F_ROH_ was 0.29 ± 0.067 (standard deviation), 0.30 ± 0.082, 0.28 ± 0.061, and 0.27 ± 0.038 for Wild 2019, Mutant 2019, Wild 2022, and Mutant 2022 groups, respectively. The four groups did not differ significantly regarding inbreeding estimates (Table S6).

**Figure 3.**
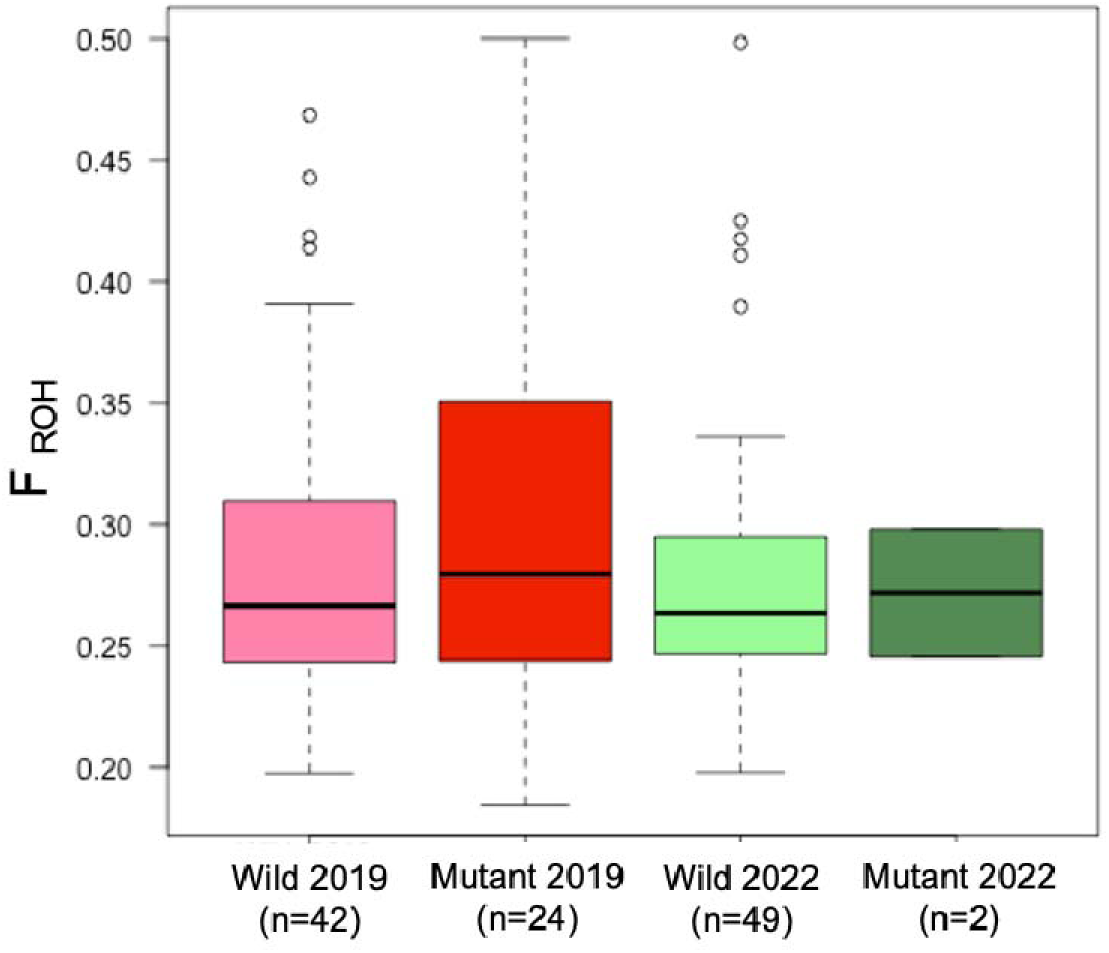
Inbreeding estimates based on F_ROH_ and genome-wide SNP data. No significant difference was observed between populations (Welch’s t-test, *P* > 0.05). We analyzed the genetic structure through clustering using ADMIXTURE (Alexander et al. 2009) and principal component analysis (PCA). Clustering results showed that the cross-validation error was lowest when K = 3 (Fig. S3), with no obvious structure in the model with K = 3 (Fig. 4A). Likewise, PCA did not identify any obvious components that explained variation in the dogs (Fig. 4B).

We applied the neighbor-joining method for each PWC and Nei’s genetic distance to infer genetic relationships between groups. The phylogenetic tree revealed one clade that included all Mutant dogs and another clade including all Wild dogs (Fig. 4C). The population-based Nei’s genetic distance indicated genetic similarities within the Mutant groups and within the Wild groups (Fig. 4D). Both analyses revealed that PWCs homozygous for the *SOD1* mutation were more common in certain lineages.

**Figure 4.**
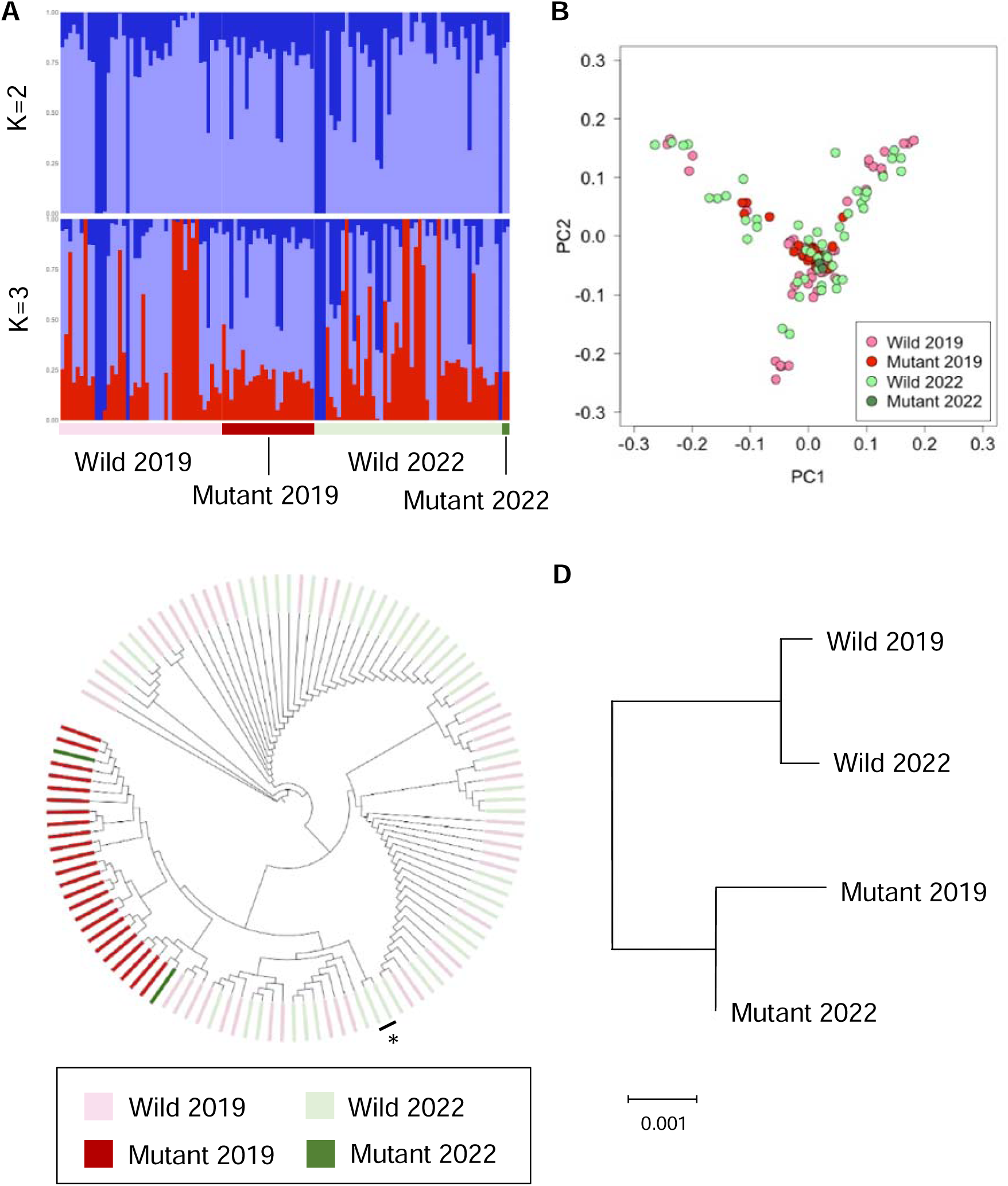
Genetic structure and relatedness of PWCs. A: ADMIXTURE. B: PCA. C: Neighbor-joining phylogenetic tree. The asterisk of a lineage includes only PWCs tested in 2022, indicating the dogs from different lineages in the 2019 group. D: Dendrogram based on Nei’s genetic distance.

To estimate the potential number of dogs bred, we used the linkage disequilibrium method to determine the contemporary *N*e (Do et al. 2014). The contemporary *N*e of Wild 2019 (n = 42) was 48.9, lower than that of Wild 2022 (73.1, n = 49).

## Discussion

Our analysis revealed that the availability of DTC genetic testing coincided with a decrease in the frequency of homozygous *SOD1*: c.118G<A mutation among PWCs, reflecting negative selection against the mutation. Our study provides valuable empirical evidence that genetic testing coupled with selective breeding can lower mutation frequency in the span of a few years. With the widespread, global availability of commercial genetic testing for *SOD1* mutations (Neeves and Granger 2015), breeding programs can apply the test results and make informed systematic mate selection decisions to decrease the frequency of this deleterious variant.

Genetic testing for adults and puppies started in 2017 and 2019, respectively. In this study, the simulation, encompassing both large and small models, revealed a significant decrease in the allele frequency of the mutation between 2017 and 2018. In addition, *F*_ST_ analysis and the simulations suggested a strong selective pressure on the mutation between 2019 and 2022. These results indicate that genetic testing for adults and puppies has led to a decrease in the frequency of the mutation. This decrease could be attributed to the introduction of large-scale genetic testing for dogs by Japanese pet shops and breeders since 2017. Subsequently, breeders may have avoided mating parents carrying the mutation. This prevented the production of puppies with the mutation and continuously reduced its frequency between 2017 and 2022.

Studies investigating selection signatures during dog domestication (Akey et al. 2010; Wang et al. 2013; Plassais et al. 2019) have identified an influence on phenotypes such as body size, coat color, and behavior. To date, selection scans have mainly focused on a given region over 10,000 years or similarly long periods (Akey et al. 2010). In contrast, selection scans over short periods (a few years) in mammals are rare. Therefore, our results provide insights into the genome evolution of mammals.

EHH is a widely used statistic in genome biology and evolutionary genetics to detect regions of recent or ongoing positive selection (Sabeti et al. 2002). This statistic quantifies a haplotype that quickly sweeps toward fixation, making it effective for detecting hard selective sweeps. Theoretically, this observation can be detected in populations with random mating; however, our study revealed longer haplotypes in the *SOD1* region in the 2019 group, as opposed to the 2022 group, where a strong selection occurred. A potential reason for the shorter haplotypes in the 2022 group could be genotype-based selection, employing breeds among a larger number of dogs from multiple lineages. First, given that PWCs are inbred breeds, the longer haplotypes observed in the 2019 group could be considered normal and not indicative of selection. Over the course of three years, our SNP-based *Ne* estimation suggests that approximately 1.5 times more dogs have been introduced and bred. Different lineages are employed during mating to prevent pairing with dogs carrying the DM mutation, a practice substantiated by our genetic analyses. Based on the genetic testing results, breeders mate dogs without the mutation, leading to a high number of recombinations around the mutation site. Future research, potentially employing model-based and/or observation-based approaches, will be required to address this possibility.

While previous studies have focused on phenotype-driven selection, our current study focuses on genotype-driven selection. Our results indicate that selective breeding was based on the *SOD1* mutation, which causes canine DM. Here, in addition to *SOD1*, we identified three candidate genes for selection (*ABCA4*, *TNXB*, and *COL11A2*) that have been reported to be associated with disease phenotypes. *ABCA4* is associated with an autosomal recessive retinal degenerative disease in Labrador retrievers (Mäkeläinen et al. 2019). Whole-genome sequencing of mixed-breed dogs has revealed two missense variants in *TNXB,* which caused an Ehlers-Danlos syndrome-like signature (Bauer et al. 2019). Additionally, the missense variant of *COL11A2* is associated with skeletal dysplasia 2 and is inherited as a monogenic autosomal recessive trait with incomplete penetrance, primarily in Labrador retrievers (Frischknecht et al. 2013). These genes may be associated with *SOD1* or DM while also being linked to valuable traits beneficial for individual survival and/or breeding performance. Since our study revealed selection signatures based on *F*_ST_ and genetic drift simulations, further research is essential to identify selective pressure and the genes involved. This study also had a limitation. Unknown admixture from other lineages may have introduced threshold bias, specifically for simulation-based thresholds. For example, we detected a migration signature in the minor alleles of two SNPs (BICF2G630168514 and BICF2G630168527, see Table S3), which were detected in the 2022 group but not in the 2019 group. This might be attributed to admixture from other lineages identified in the 2022 group, as indicated by our population and phylogenetic analyses. The simulation model employed here assumed a constant *Ne*, although our results indicate that breeding size possibly increased between 2019 and 2022. Therefore, unknown admixture events and fluctuations in breeding size were not factored in, potentially leading to an underestimation of the threshold.

The MHC plays a central role in pathogen resistance (Debenham et al. 2005). Canine MHC, also referred to as dog leukocyte antigen, exhibits genetic variations (Kennedy et al. 2007; Niskanen et al. 2013) that are associated with autoimmune diseases (Jokinen et al. 2011; Gershony et al. 2019). In this study, we found two GO terms related to the MHC for candidate SNPs for selection. Considering that these terms were identified from the test results of pups, our findings imply that selective pressure on the MHC region was related to survival in pups but not in adults, possibly because the kennel environment necessitated the rapid development of a strong pathogen-resistant autoimmune system. Evidence of post-copulatory selection at MHC-related loci in puppies supports this hypothesis (Niskanen et al. 2016). However, we did not sequence MHC haplotypes or obtain relevant phenotypes for each dog (e.g., survival rate). Thus, further research is required to identify the mechanisms (e.g., targeting haplotypes) underlying MHC selection.

Our research also demonstrated that the mutation can be selected against without lowering the *Ne* (i.e., generating inbred animals). Inbreeding avoidance is essential for effective animal breeding (Sams and Boyko 2019). Our genetic analysis revealed no differences in Ho and inbreeding levels between the 2019 and 2022 groups. In addition, we estimated a larger *Ne* for the 2022 group (73.1) than for the 2019 group (48.9). The phylogenetic analysis (Fig. 4C) further implied that the genetic origin of some Wild dogs in 2022 was from other lineages. Taken together, our findings suggest that PWC breeders used genetic testing results to limit inbreeding while mating dogs from different families or lineages.

Notably, our study focused on the genotype of the *SOD1* mutation rather than the phenotype. The median onset of DM in PWCs is 11 years (Coates et al. 2007). Considering the start of DTC genetic testing in PWCs in Japan, the corresponding decreases in the prevalence of *SOD1*-associated disease should be noticeable around the 2030s. Accordingly, further research using comprehensive phenotypic datasets, such as those available from pet insurance companies, is required to monitor DM onset in PWCs.

In conclusion, this study highlights the value of genetic testing as a tool to lower the risk of canine DM while avoiding animal inbreeding. We conducted a genome-wide analysis of short-term selection in PWCs and found that only a few years were required to reduce the number of dogs homozygous for the mutant allele. Our results highlight that genetic testing could reduce the prevalence of predictable genetic conditions, thus contributing to improved animal welfare.

## Materials and Methods

### Ethics statement

We obtained all swab samples from dogs with the consent of their owners. The ethics committee at the Anicom Specialty Medical Institute approved the study procedures (ID:2022-01).

### Genotyping for the SOD1:c.118G<A variant and statistical testing

Buccal swabs from 5,512 PWCs were sampled by breeders/owners and pet stores from all over Japan between January 1, 2017, and December 31, 2022, with their consent. DNA was extracted from oral mucosal tissue using a commercial kit (chemagic™ DNA Buccal Swab Kit, PerkinElmer and DNAdvance Kit, Beckman Coulter). We performed real-time PCR to determine genotypes of DM-associated mutations (*SOD1*:c.118G<A), specifically wild-type homozygotes (G/G), heterozygous carriers (G/A), and variant homozygotes (A/A), as previously reported (Zeng et al. 2014).

We used Fisher’s exact test in R to compare the decrease in diploid genotypes between 2016 and 2022 (RCore 2016). Next, we employed the DriftSimulator.R in R to simulate allele frequencies under genetic drift (https://www.uni-goettingen.de/de/software/613074.html). Given the commencement of large-scale genetic testing in 2017, we used data from the previous year (2016) for the simulation. Considering the difficulties in accurately inferring *Ne*, two genetic models were considered in this study. The first, a large model, was based on the understanding that approximately 10% of dogs sold annually in Japan are used for breeding (https://www.env.go.jp/nature/dobutsu/aigo/2_data/pamph/rep_h1503.html). Thus, 540 dogs were set as the *Ne* for 2016, given that 5,395 PWCs had been registered with the Japan Kennel Club (https://www.jkc.or.jp/archives/enrollment/4598). The second, a small model, was estimated based on the SNPs in this study (see Estimating *Ne* section) and was set at 49. The breeding cycle was set as one per year, indicating that six breeding generations had passed over six years (2016–2022). The sex ratio was set at 20% for males and 80% for females. Using these parameters, we conducted 10,000 simulations of genetic drift (simulated allele frequency) and adopted the resultant distribution as the null for subsequent comparisons with observed allele frequencies through six years. Next, we calculated the 95% confidence interval of the observed allele frequency based on a binomial distribution using the *binom.test* function implemented in R. Furthermore, we calculated the probability of the observed allele frequency being lower than that of the simulated allele frequencies. The probability was determined by dividing the number of simulated allele frequencies lower than the observed frequency by 10,000. Significance was set at *P* < 0.05 for both tests.

### Genome-wide SNP genotyping

We used an SNP-genotyping array (Canine 230 K Consortium BeadChip Array, Illumina, San Diego, CA, USA) to investigate the effects of genetic testing on the whole genome. We compared genome-wide SNPs from four subpopulations of wild-type and variant homozygotes of the *SOD1* risk allele, all derived from dogs born in 2019 and 2022. Considering the effect of genetic background on array-based analyses, we selected all dogs from the two largest clients for array-based analyses (Table S1). We used an SNP-genotyping array (Canine 230 K consortium) and the Illumina iScan system to detect over 230 K SNP genotypes. Genotype coordinates corresponded to the CanFam3.1 genome assembly.

We conducted quality control using PLINK version 1.90 (Chang et al. 2015). Missing rates for each locus (geno option) and individual dogs (mind option) were both < 1%. We did not examine the Hardy-Weinberg equilibrium as we did not assume random mating for pedigreed PWC. Population genetic analyses should not include close relatives; therefore, we conducted robust relationship-based pruning (Manichaikul et al. 2010) using the king-cutoff option in PLINK version 2.00a2LM, with a threshold of 0.176 to exclude pairings between monozygotic twins and first-degree relatives. For allele frequency-based quality control, we excluded SNPs with a minor allele frequency of < 0.01. We also removed sex chromosome variants and autosomal indels (insertions-deletions) because their effects on allele frequency differ from those of autosomal SNPs. The final analysis retained 143,013 SNPs. After quality control for individual dogs, including the removal of related animals, 117 dogs remained: homozygous wild-type (G/G) dogs tested in 2019 (Wild 2019, n = 42) and in 2022 (Wild 2022, n = 49) as well as homozygous mutant dogs tested in 2019 (Mutant 2019, n = 24) and in 2022 (Mutant 2022, n = 2; Table S2). We employed this dataset for subsequent analyses.

### Detecting selection pressure on SNPs

Artificial selection alters the frequency of genes associated with phenotypes. To identify selection signatures in the PWC genome, we estimated *F*_ST_ in the 2019 and 2022 groups. We estimated *F*_ST_ values using the –fst option implemented in PLINK version 1.9. The first threshold for selective pressure was based on SNPs with *F*_ST_ values in the upper 0.1%. In addition to the *F*_ST_-based scan, we conducted a more rigorous detection using a simulation-based scan. We also implemented a simulation based on genetic drift, conducting it 10,000 times. The small model (*Ne* = 49) was used to assign a rank, indicating the probability of frequency to increase/decrease in 2022 compared to that in 2019. Subsequently, we set the second threshold to below 0.025 (rank 250) or above 0.975 (rank 9750; two-sided 2.5 percentile) as the suggestive level and performed additional Bonferroni correction for 143 SNPs below 0.001 (rank 1) or above 0.999 (rank 9999; two-sided 1.7 percentile) as the significant level. The SNPs that met the criteria of the first threshold were subjected to the simulation-based scan. The SNPs that passed the suggestive level were denoted as “candidate SNPs for selection” and those that passed the significant level were denoted as “selected SNPs”. Genes in the ± 100 kb region surrounding SNPs which passed thresholds with suggestive and significant level were considered candidate selected genes.

Furthermore, to investigate haplotype length surrounding *SOD1* mutation, we calculated the EHH of wild-type populations. The EHH is a measure used to detect genome regions under recent selective pressure (Sabeti et al. 2002). First, target SNPs under selective pressure in the Wild 2019 and 2022 populations were phased using Beagle 5.4 (version 22Jul22.46e) (Browning et al. 2018) with default settings. After phasing, EHH for the target SNP was estimated using a 1 Mbp window in Selscan version 2.0.0 (Szpiech and Hernandez 2014).

To determine the functional classes of genes associated with the analyzed traits, we performed a GO analysis powered by PANTHER (Thomas et al. 2022). We selected target genes using linkage disequilibrium analysis and the PANTHER overrepresentation test (release 20221013). ‘Molecular function’ was used for domestic dog (*Canis lupus familiaris*) dataset annotation in PANTHER 17.0. Ensembl gene IDs were used for data annotation. Statistical differences were assessed using Fisher’s exact test. A false discovery rate of *P* < 0.05 was considered significant.

We obtained all gene datasets from the Ensembl Genome Browser (release 104; CanFam 3.1) and retrieved Ensembl gene IDs using Bedtools version 2.27.1 (Quinlan and Hall 2010; Danecek et al. 2021).

### Inbreeding levels

We inferred inbreeding levels using specimens from genome-wide SNP genotyping. The R package DetectRuns was used to obtain the proportion of times each SNP fell inside a run per population, corresponding to locus homozygosity or heterozygosity in the respective population. We used the following DetectRuns parameters: minSNP = 41, maxGap 10^6^, minLengthBps = 50000, and minDensity = 1/5000.

### Population genetics

To clarify the genetic structure and phylogenetic relationships in the PWC population, we performed PCA using PLINK version 1.9 with default settings and maximum-likelihood ancestry analysis using ADMIXTURE version 1.3 (Alexander et al. 2009). For ADMIXTURE, we set the number of populations (K) between 2 and 10. We used the option cv for cross-validation error calculation to select the optimal K value based on the lowest error.

For genetic relationships, we first converted the PLINK PED format into FASTA format. We then converted the FASTA file to NEXUS format in MEGA X (Kumar et al. 2018). We constructed a phylogenetic tree using the neighbor-joining algorithm with p-distance in MEGA X.

We calculated Nei’s standard genetic distance D (Nei 1987) between the four populations (Wild 2019, Wild 2022, Mutant 2019, and Mutant 2022). Nei’s D was estimated using GenoDive version 3.06 with default settings (Meirmans and Van Amsterdam 2019).

### Estimating Ne

The contemporary *N*e of PWC was estimated from genome-wide SNP data using a linkage disequilibrium method in NeEstimator version 2.1 (Do et al. 2014). The lowest allele frequency was set at 0.01.

## Data availability

SNP data for the PWCs are available from the Dryad database (DOI: 10.5061/dryad.rbnzs7hhk)

## Acknowledgments

We thank all participating breeders, veterinarians, and dog owners for collecting buccal swabs from their dogs. We also thank Dr. Ryo Horie, Mr. Ryuta Kuwana, and all the Anicom Specialty Medical Institute members for their feedback on this study. This study was supported by the Research Foundation of Anicom Specialty Medical Institute.

## Author Contributions

YM and HU designed the study; HU, NA, FY, GI, KO, and YM collected dog samples, performed experiments, contributed to the analytical tools, analyzed the data, and wrote the paper.

